# METTL3 Modulates Radiation-Induced Cardiac Fibrosis via the Akt/mTOR Pathway

**DOI:** 10.1101/2024.09.07.611836

**Authors:** Shunsong Qiao, Chao Tang, Jingjing Zhu, Yu Feng, Li Xiang, Jing Zhu, Xiaosong Gu

**Affiliations:** Department of Cardiology, the second affiliated hospital of Soochow University, 215000 Suzhou, Jiangsu, China; Department of Endocrinology, the second affiliated hospital of Soochow University, 215000 Suzhou, Jiangsu, China

**Keywords:** Radiation damage, Myocardial fibrosis, m6A, METTL3, Akt signaling

## Abstract

**BACKGROUND:** Radiation therapy for cancer treatment frequently leads to radiation-induced heart disease (RIHD), characterized as cardiac fibrosis and heart failure. The enzyme METTL3, a key player in RNA methylation, is known to influence various cellular processes, while its specific role and mechanism in RIHD are not well understood.

**OBJECTIVES:** The purpose of this study was to explore the potential mechanism of METTL3 in regulating cardiac fibrosis induced by radiation.

**METHODS:** We constructed an X-ray-modulated RIHD mice models to investigate the role of METTL3 in cardiac fibroblasts. In parallel, METTL3 overexpression and silence were conducted on fibroblasts and mice heart to evaluate pro-fibrotic protein expression, cardiac fibrosis, and heart function.

**RESULTS:** Elevated METTL3 expression was observed in both irradiated cardiac tissues and fibroblasts. Moreover, overexpression of METTL3 in cardiac fibroblasts was associated with increased expression of pro-fibrotic proteins and exacerbated fibrosis, whereas METTL3 silencing attenuated these adverse effects. Likewise, METTL3 knockdown ameliorated cardiac dysfunction and reduced fibrosis. Taken together, METTL3 could induce cardiac fibrosis via activating the Akt/mTOR signaling pathway to promote fibroblast proliferation and myofibroblast differentiation.

**CONCLUSIONS:** METTL3 is a critical of RICD through modulating the Akt/mTOR pathway, hence targeting METTL3 presents a promising therapeutic strategy to mitigate the adverse cardiac effects of radiation therapy in cancer patients.

## 1. Introduction

Radiation therapy has emerged as an important treatment for cancer. A number of research evidence have suggested a cause-and-effect relationship between cardiovascular disease and mediastinal radiotherapy. Radiation therapy is a common cause of heart problems^[1]^. The heart is usually exposed to radiation when the patient undergoes mediastinal radiotherapy. As a consequence, radiation-induced heart damage or radiation-induced heart disease (RIHD), which comprises a spectrum of cardiac conditions including pericardial disease, myocardial infarction, myocardial fibrosis, coronary artery disease, and heart failure, can develop ^[2,3]^. It is estimated that 14% of patients who are exposed to 1 Gy of radiation will develop RIHD^[4]^. Moreover, as the mean cardiac dose escalates, the incidence of major cardiac events increases by 7.4% annually, and the adverse outcome usually occurs within 5-20 years post-radiotherapy in women who survived breast cancer^[1]^.

Tissue fibrosis is an important feature of RIHD, cardiac fibroblasts (CF) play a crucial role in remodeling the external environment via synthesis and degradation of ECM^[5]^, particularly during. cardiac repair, thereby maintaining cardiac homeostasis. Upon exposure to radiation, multiple signaling pathways could coordinate and activate at genetical, molecular and cellular levels in CF for both short and long-term, thereby promoting transformation into myofibroblasts. Subsequently, excessive extracellular matrix (ECM) and collagen released from the myofibroblasts can ultimately contribute to myocardial fibrosis^[6]^.

The N6-methyladenosine (m6A) modification of RNA is a prevalent and highly conserved across mammalian cells, crucial for maintaining the stability of RNA, controlling translation, and facilitating other intracellular biological processes^[7]^. The main catalytic subunit in the methyltransferase complex is methyltransferase like 3(METTL3), regulates numerous biological processes^[8]^. METTL3 could serve as a biomarker for the diagnosis or treating cardiovascular diseases^[9]^, given its role in pressure overload, ischemia/reperfusion injury, adverse cardiac remodeling post-myocardial infarction. Recent studies indicate that m6A is closely associated with the DNA damage response (DDR)^[10]^. At present, it is thought that RIHD is a result of various mechanisms interacting with each other through multiple complex pathways, DNA damage response, oxidative stress (OS) and inflammation, endoplasmic reticulum and mitochondrial damage are considered to be the main reasons^[4]^. And it has been suggested that DNA-related damage is a critical factor in the development of RIHD^[11]^. However, the specific role of METTL3 in RIHD remains unclear.

Previous research has identified the PI3K/AKT signaling pathway as playing a pivotal role in the development of acute radiation proctitis^[12]^. Furthermore, researchers have discovered that elevated levels of METTL3 and hyperactive m6A modifications are associated with pulmonary fibrosis. In the context of pulmonary fibrosis, m6A modifications contribute the onset of pulmonary fibrosis by activating the PI3K-AKT-mTOR signaling pathway^[13]^. We hypothesize that there may be a significant interrelationship among METTL3, Akt, and RIHD, although the underlying mechanisms remain to be elucidated. As a result, our study aims to assess the role of the methyltransferase METTL3 in radiation-induced cardiac fibrosis, as well as investigate the mechanisms by which METTL3 influences the Akt pathway in the context of radiation-induced cardiac fibrosis.

## 2. Methods

### 2.1. Animals

The C57BL/6 mice were purchased from Nanjing Jicui Pharmachem Biotechnology Co. All animal protocols adhered to the guidelines established by the Institutional Animal Care and Use Committee and the Ethic Committee of Soochow University.

### 2.2. Primary cardiac fibroblasts

Cardiac fibroblasts were harvested from 1∼3-day-old neonatal mice. Hearts were rapidly excised and immersed in DMEM medium (10-013-cv, Corning, USA) containing 2% double antibody pre-cooled, well sheared (less than 1 mm^3^), and digested in 0.25% trypsin (Thermo Fisher, USA) and 0.1% collagenase II (BioFroxx, Germany) at 37°C. After digestion, cells were collected, centrifuged at 1000 rpm for 3 minutes, and then cultured in DMEM supplemented with 10% fetal bovine serum (Thermo Fisher, USA), 100 U/ml penicillin, and 100 μg/ml streptomycin (Thermo Fisher, USA) for 2 h. The medium was then changed for the first time to remove weakly adherent cells. Isolated purified fibroblasts were further incubated in DMEM at 37°C in humidified air with 5% CO2 and 95% O2. Routine passages after 2 to 3 days, P1-2 were used for experiments.

### 2.3. Irradiation

Control group, 10Gy irradiation group, and 20Gy irradiation group, with the division being done randomly. Subsequently, the mice underwent a single irradiation session and were fed for a period of 28 days. Irradiation was performed using a single-field technique focused on the chest. The irradiation field had dimensions of 1×1 cm^2^ and a depth of 1.5 cm. The source skin distance was set at 100 cm. The irradiation volume was 10Gy or 20Gy. The irradiation was delivered using a linear accelerator (Elekta Synergy, UK) with a dose rate of 600 cGy/min. The energy of the radiation used was 6 MV X-ray^[14]^. The hearts were collected for histologic analysis 4 weeks after single-dose irradiation.

Cardiac fibroblasts were divided into five groups and irradiated with 0, 2, 5, 10 and 20 Gy X-rays. Cells were irradiated with a medical linear accelerator (Elekta Synergy, UK) with a beam energy of 4 MV x-rays and a dose rate of 600 cGy/min; the source surface distance (SSD) was 100 cm. Primary cardiac fibroblasts were seeded in culture plates overnight before exposure to incremental doses of ionizing radiation (0, 2, 5, 10, and 20 Gy)^[15]^.

### 2.4. Bioinformatics studies

The post-radiation cardiac injury RNA-seq dataset was retrieved from the GEO database (https://www.ncbi.nlm.nih.gov/geo/), and the dataset: GSE218447 was selected and downloaded. For pathway enrichment analysis, the “clusterProfiler” R package v 3.16.1 Kyoto Encyclopedia of Genomes (KEGG) was analyzed. Pathways with *P*-value less than 0.05 were considered significantly enriched. Selected significant pathways were plotted as bar graphs.

### 2.5. m6A Dot Blot Assay

The mouse hearts and CFs RNA was extracted using TRIzol reagent (Thermo Fisher Scientific, USA) as per the manufacturer’s instructions. RNA concentration was measured using a NanoDrop ND-1000 spectrophotometer (Agilent). m6A dot blotting experiments are shown below. Briefly, RNA samples were loaded onto Amersham HybondN+ membranes (Biyun Tian, FFN13) and UV crosslinked. The membranes were blocked with 5% skim milk for 1 h, incubated with m6A antibody (Epigentek, P-9005-48, 1:5000) overnight at 4°C, and left with horseradish peroxidase (HRP)-coupled secondary antibody (Santa Cruz Biotechnology; Sc-2030) for 1 h at room temperature. The membranes were visualized using an enhanced chemiluminescence (ECL) detection system (Amersham Biosciences, Piscataway, NJ). The relative signal density at each point was quantified using Image J software.

### 2.6. Cardiac METTL3 knockdown and overexpression

Cardiac METTL3 knockdown mice model was established by injecting knockdown METTL3 and negative control virus into the left ventricular myocardium. One week later, both groups and the non-viral transfected group were irradiated simultaneously, and the effect was assessed after four weeks.

METTL3-specific shRNA (shMETTL3) and a negative control (sh-scr) were synthesized by OBiO Technology (Shanghai, China). Transfection with shMETTL3 led to decreased METTL3 expression. Transfection of shRNA also induced knockdown of other target genes. Gene silencing was achieved by transfection of a pre-designed shRNA duplex designed and synthesized by OBiO Technology (Shanghai, China). CFs were seeded in six-well plates at 50–60% confluency and treated with METTL3 shRNA lentiviral particles. 24h later, cells were cultured in a complete culture medium containing purromycin (1.0μg/mL) for 12–14d. Western blotting analysis confirmed a more than 95% METTL3 knockdown in stable cell lines. The target sequence of the knockdown virus was 5’-GCACCCGCAAGATTGAGTTAT-3’ (sh-METTL3), and that of the overexpression virus was 5’-CGCAAATGGGGCGGTAGGCGTG-3’ (OE-METTL3).

### 2.7. Mouse echocardiography

Mice were anesthetized with 3% isoflurane during induction and 2% isoflurane during maintenance, and were placed on a heated carrier table to 37℃. Images were acquired using an VERMON probe (23 MHz) on a VINNO6 Dimension color Doppler ultrasound machine. Analysis was conducted using the EchoPac workstation and 2D strain imaging software. The optimal and distinct images were acquired by depilating the precordial region of mice and capturing the short-axis view of the left ventricular papillary muscle at the sternum, as well as the apical 4-chamber view. The M-mode ultrasound was used to assess the LV end-diastolic internal diameters (LVIDd) and end-systolic internal diameters (LVIDs) in the parasternal long-axis view of the left ventricle. Subsequently, ejection fraction (LVEF) and left ventricular shortening rate (LVFS) were calculated. The images were then imported into the EchoPac workstation and the researchers analyzing the echocardiograms were unaware of the animal genotype and/or intervention at the time of analysis.

### 2.8. Wheat germ agglutinin (WGA) staining

After dewaxing and rehydration, the heart slides were incubated with Alexa Fluor 488 (Invitrogen, W11261, 5 μg/ml) coupled to WGA for 10 min at room temperature. To quantify cardiomyocyte size, five individual hearts per group (at least 100 cells) were captured near the apex with a laser-scanning confocal microscope (LSM 700, Zeiss). The size of each cell was quantified using Image J software.

### 2.9. Tissue histological analysis

Hearts were collected 4 weeks after irradiation, fixed in 4% paraformaldehyde overnight, embedded in paraffin, and sliced into 5μm-thick cross sections starting at the apex. Heart paraffin sections were baked in an oven at 60℃ for 60 minutes and then deparaffinized with xylene. After hydration in graded alcohol and water, the sections were stained with Masson Trichrome kit (Solarbio, Beijing, China). In these sections, fibrotic tissue was blue and myocardial tissue was red. The percentage of fibrotic tissue, blue collagen deposition on Masson trichrome-stained sections, was measured using Image-Pro Plus 6.0. Heart sections were stained with Sirius Red kit (Solarbio, Beijing, China), and fibrotic tissue was red and myocardial tissue was yellow. The collagen area was calculated using Image J software.

Hearts were embedded in paraffin, sectioned into 5 μm slices, deparaffinized, rehydrated, and antigen retrieval. Sections were permeabilized with 0.5% Triton X-100/phosphate buffer saline, then blocked with 5% goat serum (Jackson ImmunoResearch Laboratories, USA) for 1 hour at room temperature and incubated with primary antibodies overnight at 4°C. After PBS washing, the slices were incubated with the corresponding fluorescence-coupled secondary antibody for 1 hour at room temperature, followed by DAPI (Sigma) counterstaining. Primary antibody used were α-SMA (abcam, ab7817, 1:100) and Vimentin (proteintech, 60330-1-lg, 1:100). Secondary antibodies: goat anti-mouse Alexa Fluor 488 (Thermo Fisher, A-11001, 1:500); goat anti-mouse Alexa Fluor Plus 594 (Thermo Fisher, A32742, 1:500).

### 2.10. Cell proliferation and migration assays

CFs cells were seeded into six-well plates (2 × 10^5^ cells in each well) and cultured for 24 hours. Cell proliferation was quantitatively measured using the EdU Apollo-567 assay kit (RiboBio, Guangzhou, China). Cell nuclei were stained with EdU and DAPI(4,6-diamino-2-phenyl indole) and observed under a fluorescence microscope (Leica, DM 4000, Germany). Cells in each field of view were then counted and analyzed. The nuclear EdU ratio,% with DAPI, was calculated for at least 1,000 cells in five randomized views per treatment.

CFs cells (40,000 cells per chamber in serum-free medium) were added to the upper Transwell chambers (8 μm wells, Corning, New York, NY, USA), and the lower chambers were filled with 10% FBS(fatal bovine serun) complete medium. 24 hours later, CF cells migrating to the lower chambers were fixed, stained, and counted.cataway, NJ).

### 2.11. Western blot analysis

Protein extraction from mouse heart tissue and cardiac fibroblasts. Aliquots of 30 μg of protein from each sample were separated by 10% SDS-polyacrylamide gel electrophoresis (SDS-PAGE) (processed as shown in the legend) and transferred to polyvinylidene difluoride (PVDF) membranes (Millipore, Bedford, MA). The membranes were blocked with 10% skim milk powder for 1 hour, incubated with the specific antibody overnight at 4°C overnight, and the secondary antibody (HRP-coupled anti-rabbit or anti-mouse IgG, at an appropriate dilution) for 45 minutes to 1 hour at room temperature. Antibody binding was detected using an enhanced chemiluminescence (ECL) detection system (Amersham Biosciences, Piscataway, NJ). The primary antibodies used were as follows: anti-Col I (Abcam, ab21286), anti-METTL3 (Cell Signaling Technology, 86132S), anti-GAPDH (Proteintech, 10028230), anti-TGF-β1 (Proteintech,21898-1-AP),Phospho-Akt(CST,#13038);Akt(SantaCruzBiotechnology,# E2121),Phospho-PI3K(CST,#17366);PI3K(CST,#4249),Phospho-mTOR(CST,#5536); mTOR(CST,#2983),Phospho-S6(CST,#34475);S6(SantaCruzBiotechnology,#B2023). Membranes were examined on Odyssey infrared scanning system (LI-COR Biosciences, Lincoln, Nebraska, USA), and the Western blot images were captured with the band densities quantified using Odyssey 3.0 software.

### 2.12. Statistical analysis

Results are presented as mean ± SEM, and analyzed using one-way ANOVA with Tukey’s test for individual group comparisons. *P*-values < 0.05 were considered statistically significant. The analysis and presentation were performed using Prism 9(GraphPad, USA). The in vitro experiments were repeated at least three times and consistent results were obtained.

## 3. Results

### 3.1. Radiation impairs heart function and induces myocardial fibrosis

The heart function parameters indicated that radiation led to contractile dysfunction. Compared with the controls, both left ventricular ejection fraction (LVEF) and left ventricular fractional shortening (LVFS) decreased in mice after irradiation for 4 weeks. The radiation also increased left ventricular end-diastolic internal diameter (LVIDd) and left ventricular end-systolic internal diameter (LVIDs)(Figure 1A). These structural defects were accompanied with larger cardiomyocytes (Figure S1A). The overall cardiac dysfunctions were more pronounced at a dose of 20 Gy compared to 10 Gy. (Figure 1A and S1A). In parallel, the tissue histological analysis demonstrated that radiation induced myocardial fibrosis in adult mice. Significant tissue expressions of fibrotic markers including α-SMA, collagen I and TGF-β1 were all enhanced in adult mice heart after irradiation.

**Fig 1.**
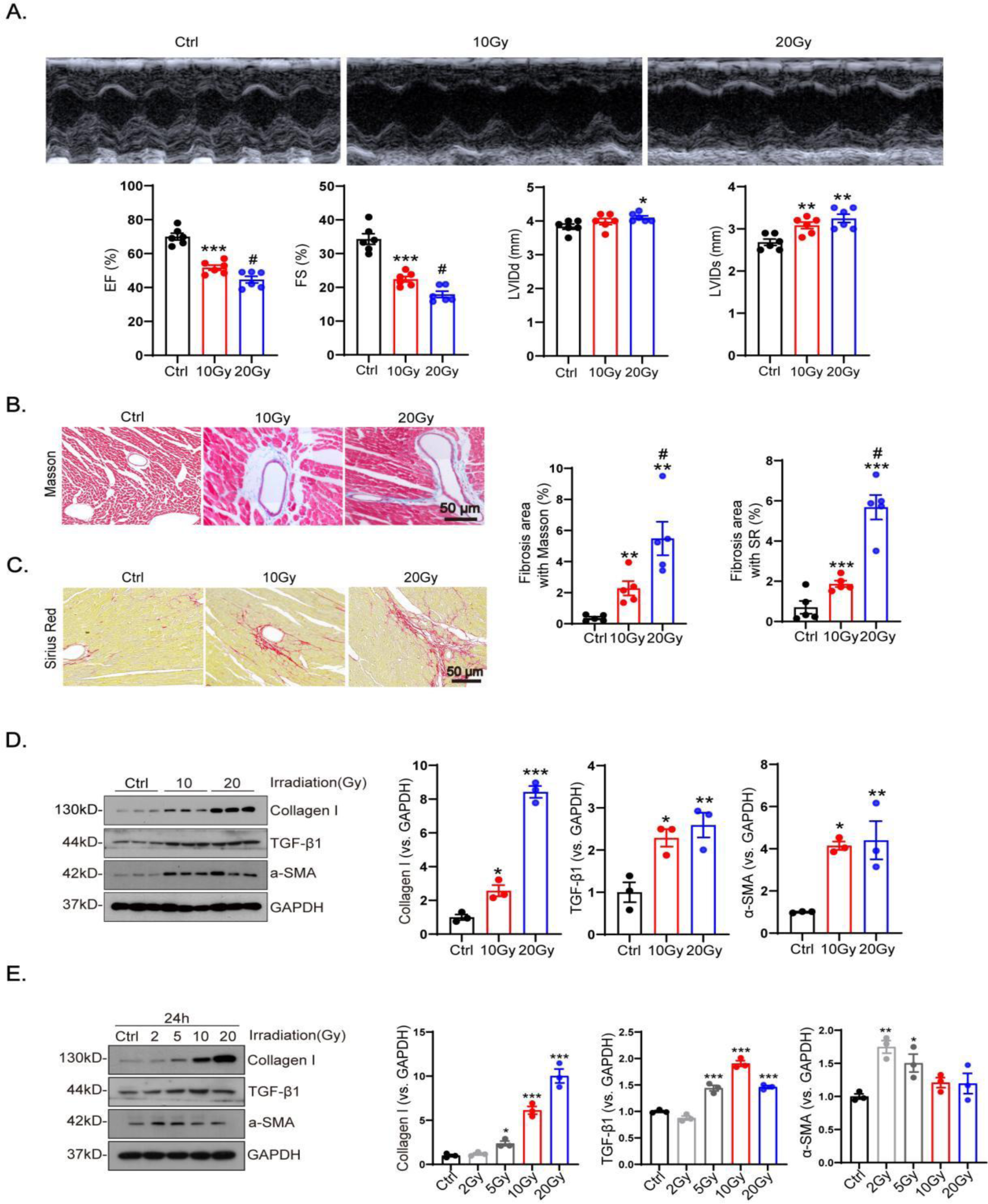
Adult mice and cardiac fibroblasts were subjected to irradiation at doses of 0Gy (Ctrl), 2Gy, 5Gy, 10Gy and 20Gy, with exposure for 4weeks and 24 hours, respectively. (A) Heart function parameters including left ventricular ejection fraction (LVEF), left ventricular fractional shortening (LVFS), left ventricular internal diameter (LVID), and left ventricular posterior wall thickness (LVPWd) were assessed using echocardiography. Tissue fibrosis was evaluated using Masson (B) and Sirius (C) red staining. Fibrotic markers including a-SMA, collagen I and TGF-β1 in heart tissues (D) and cardiac fibroblasts (E) were quantified using western blotting. Data are expressed as mean ± SEM for groups of 3, 5 or 6. All results were analyzed using one-way ANOVA with Tukey’s post hoc test and confirmed by three independent experiments. ***P<0.001, **P<0.01, *P<0.05 vs. Ctrl; ##P<0.01, #P<0.05 vs. 10Gy.

Masson, Sirius red staining of cardiac tissue sections was performed to analyze the degree of cardiac fibrosis, which showed that the degree of cardiac fibrosis increased with increasing irradiation dose, and 20Gy was more fibrotic compared to 10Gy. (Figure 1B-C). The western blot results showed that with the increase of irradiation dose, the protein expression levels of a-SMA and collagen I gradually increased in the heart tissue and CFs. Meanwhile, TGF-β1 secreted by the heart also increased with the increase of irradiation dose (Figure 1D-E). Further immunofluorescence staining of heart sections showed that perivascular fibrosis of the heart worsened with increasing irradiation dose (Figure S2A).

In addition, immunofluorescence staining of CFs showed that the fluorescence intensity of a-SMA in cardiac fibroblasts gradually increased with the increase of X-ray radiation dose, indicating an increase in the degree of fibrosis and trans differentiation of CFs into myofibroblasts. (Figure S2B).

### 3.2. Radiation induces METTL3-mediated m6A modification

A dot blot assay revealed that the irradiated group exhibited a significantly higher level of m6A modification in total RNA compared to the control group in cardiac tissues (Fig. 2A) and in CFs (Fig. 2B). In parallel, the dose-dependent increase in METTL3 levels was also demonstrated by western blot in cardiac tissues (Fig. 2C). Consistent with the findings in heart tissue, cardiac fibroblasts exposed to X-rays at varying doses (0, 2, 5, 10, and 20 Gy) showed a marked, dose-dependent increase in METTL3 expression (Fig. 2D).

**Fig 2.**
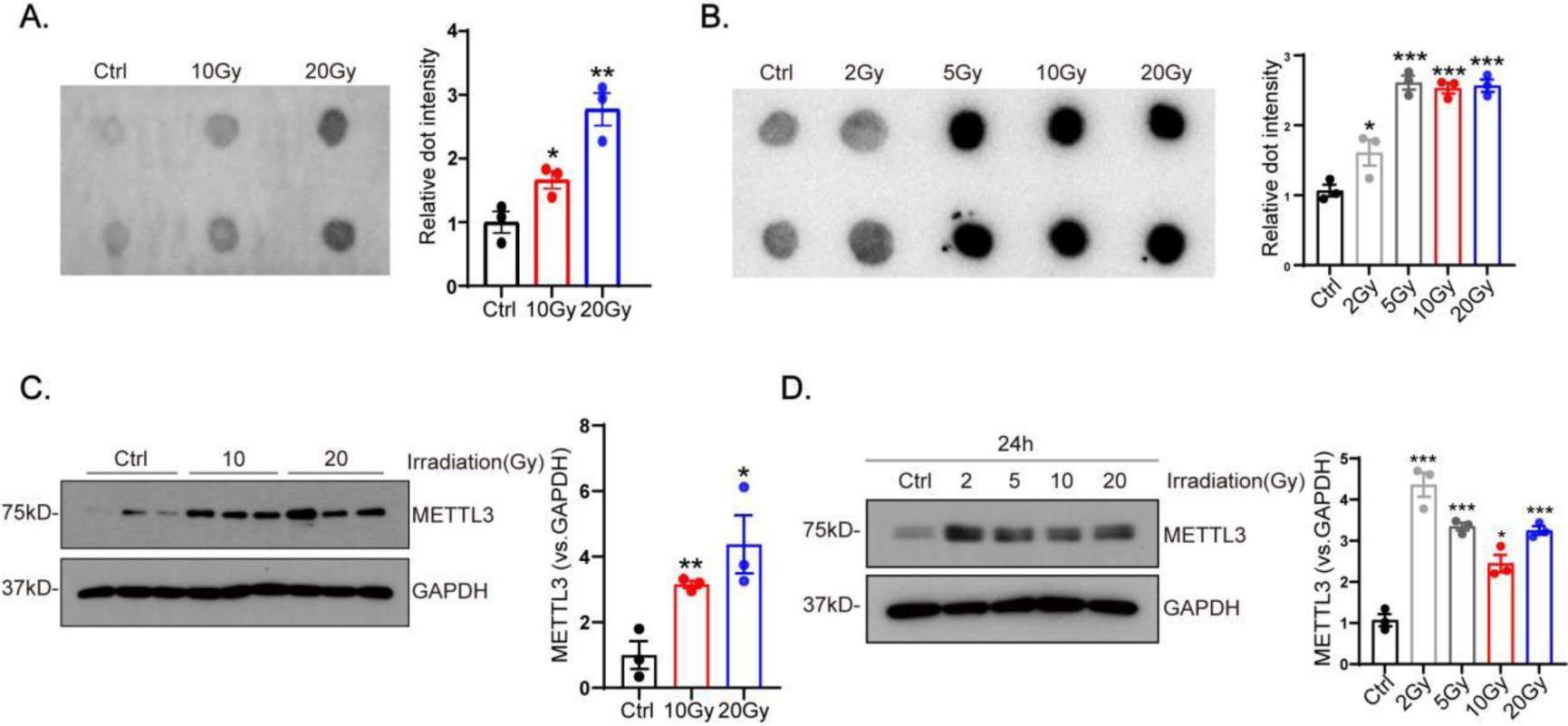
Radiation induces METTL3-mediated m6A modification of mRNA Adult mice and cardiac fibroblasts were subjected to irradiation at doses of 0Gy (Ctrl), 2Gy, 5Gy, 10Gy and 20Gy, with exposure for 4weeks and 24 hours, respectively. m6A RNA modification in heart tissues (A) and cardiac fibroblasts (B) were measured using dot blot assay. METTL3 expression in in heart tissues (C) and cardiac fibroblast (D) were also quantified using western blotting. Data are shown as mean ± SEM for groups of 3. The results were analyzed using one-way ANOVA with Tukey’s post hoc test and confirmed by three independent experiments. ***P<0.001, **P<0.01, *P<0.05 vs. Ctrl.

### 3.3. METTL3 is key to impaired heart function and myocardial fibrosis during irradiation

To investigate the effects of METTL3 inhibition on myocardial fibrosis and cardiac function following X-ray irradiation, lentiviral vectors encoding shMETTL3 were injected into mouse hearts two weeks before irradiation. Western blot analysis showed that irradiated hearts exhibited more than a 5-fold decrease in METTL3 expression compared to control hearts (Fig. 3A). Western blot results indicated that blocking METTL3 significantly decreased the levels of TGF-β1, Collagen Ⅰ and α-SMA in hearts exposed to radiation (Fig. 3A). Echocardiographic analysis was performed after four weeks, and the results showed that LVEF, FS decreased significantly and LVIDs increased after irradiation, whereas the injection of shMETTL3 maintained normal cardiac function in radiation-exposed mice with no significant change in the LVEF, FS and LVIDs values compared with the control group (Figure 3B). Interstitial fibrosis was assessed four weeks post-irradiation. Masson’s trichrome and Sirius Red (SR) staining demonstrated that METTL3 inhibition reduced interstitial and perivascular collagen deposition in irradiated myocardial tissues (Fig. 3C, D). Additionally, shMETTL3 treatment lowered the fluorescence intensity in irradiated hearts (Fig. 3E). Furthermore, cardiomyocyte size did not increase in mice injected with shMETTL3 lentivirus following irradiation, whereas cardiomyocyte volume increased in the negative virus-injected group (Fig. S3A). However, there were no significant differences in METTL3 levels, heart function, or heart volume between the irradiated and sh-scr groups. This suggests that blocking METTL3 effectively prevented myocardial fibrosis and alleviated myocardial remodeling induced by X-ray irradiation.

**Fig 3.**
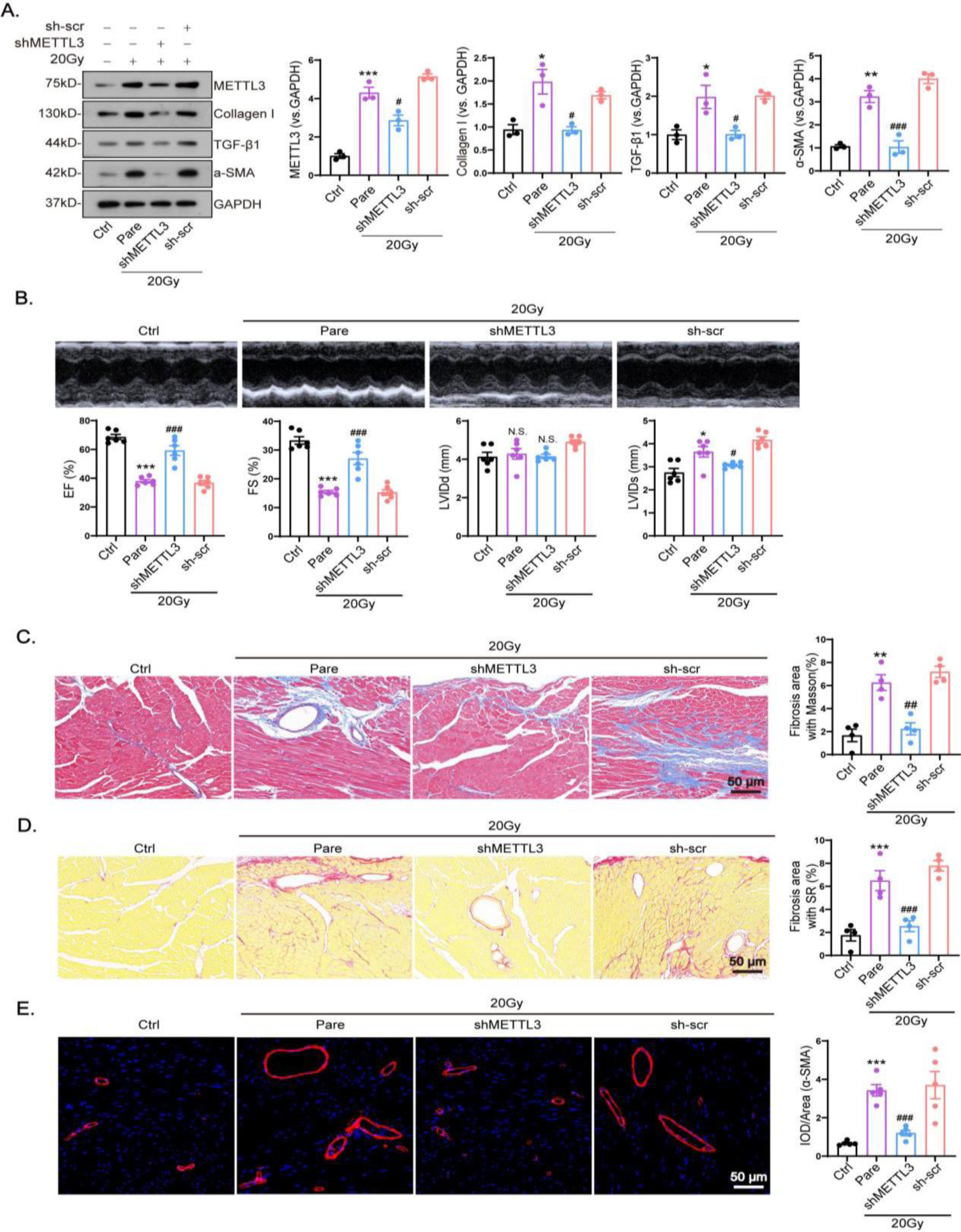
METTL3 is key to impaired heart function and myocardial fibrosis during irradiation Adult mice were subjected to irradiation at doses of 0Gy (control), 20Gy (Pare), 20Gy with cardiac co-transfection of shMETTL3(shMETTL3) or sh-vector (sh-scr) for 4 weeks. (A) Expression of METTL3 and fibrotic markers including a-SMA, collagen I and TGF-β1 in were quantified using western blotting. (B) Heart function parameters including left ventricular ejection fraction (LVEF), left ventricular fractional shortening (LVFS), left ventricular internal diameter (LVIDd), and left ventricular posterior wall thickness (LVPWd) were assessed using echocardiography. Tissue fibrosis was evaluated using Masson (C) and Sirius (D) red staining. (D) Myocardial fibrosis was evaluated using fluorescence intensity. Representative images of α-SMA(E)fluorescence intensity enhancement in irradiated hearts after METTL3 inhibition. Data are shown as mean±SEM for groups of 3,4,5 or 6. The results were analyzed using one-way ANOVA with Tukey’s post hoc test. ***P<0.001, **P<0.01, *P<0.05 vs. Ctrl; ###P<0.001, ##P<0.01, #P<0.05 vs. Pare.

### 3.4. METTL3 promotes the transdifferentiation of cardiac fibroblasts to myofibroblasts during irradiation

CFs were transfected with plasmids to reduce the expression of METTL3. Western blot result showed that the expression level of METTL3 in the knockdown group was significantly reduced compared with that in the control group (Figure S4A), and also western blot analysis showed that knockdown of METTL3 reduced the expression of a-SMA, collagen I and TGF-β1 secretion in CFs (Fig. S4A). In the EDU assay, knockdown of METTL3 was found to inhibit the proliferation of CFs (Fig. S5A), and the migration of CFs was inhibited in the transwell assay at 24 hours after transfection (Fig. S5B).

The successful overexpression of METTL3 was first verified by western blot (Fig. S4B). After that, western blot results showed that a-SMA, collagen I expression and TGF-β1 secretion were significantly up-regulated in CFs (Fig. S4B), and reduced CF proliferation in EDU experiments (Fig. S5C) and migration in permeabilized-well experiments at 24h post-transfection (Fig. S5D). Immunofluorescence analysis further confirmed that the expression of the myofibroblast marker α-SMA was increased in the METTL3 overexpression group, consistent with the results of Western blot (Fig. S5E). These findings suggest that METTL3 overexpression promotes CF proliferation, migration, and myofibroblast differentiation in vitro, while METTL3 knockdown has the opposite effect.

CFs were transfected with shMETTL3 or the corresponding negative viruses. shMETTL3 effectively decreased METTL3 expression in irradiated CFs compared to the control group (Fig. 4A). Furthermore, X-ray irradiation suppressed the expression of a-SMA, Collagen Ⅰ and the secretion of TGF-β1(Fig. 4A). X-ray irradiation significantly enhanced the proliferation and migration of CFs. However, transfection with shMETTL3 substantially reduced these irradiation-induced effects (Fig. 4B and 4C). Moreover, immunofluorescence analysis revealed that shMETTL3 treatment reduced the fluorescence intensity of CFs subsequent to X-ray irradiation (Fig. 4D), indicating that METTL3 knockdown impeded X-ray-stimulated myofibroblast differentiation.

**Fig 4.**
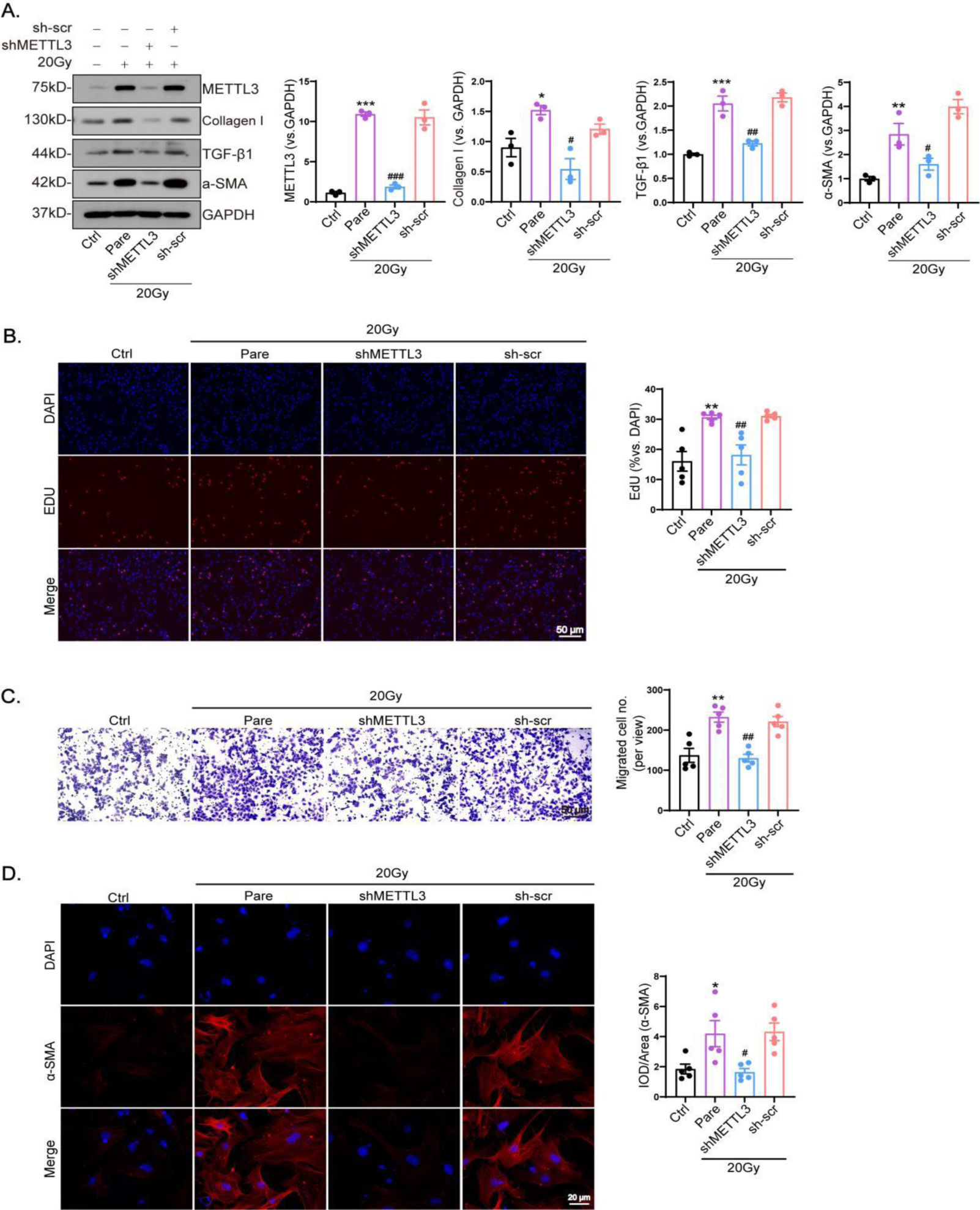
METTL3 modulates the transdifferentiation of cardiac fibroblasts into myofibroblasts during irradiation Cardiac fibroblasts were subjected to irradiation at doses of 0Gy (control), 20Gy (Pare), 20Gy with co-transfection of shMETTL3(shMETTL3) or sh-vector (sh-scr) for 24 hours. (A) Expression of METTL3 and myofibroblast markers including α-SMA, collagen I and TGF-β1 in were quantified using western blotting. Representative images and statistical analysis of shMETTL3 inhibition of radiation-induced CF proliferation (B), migration (C) Transdifferentiation to myofibroblasts (D) were also evaluated using fluorescence intensity Data are shown as mean±SEM for groups of 3 or 5. The results were analyzed using one-way ANOVA with Tukey’s post hoc test. ***P<0.001, **P<0.01, *P<0.05 vs. Ctrl; ###P<0.001, ##P<0.01, #P<0.05 vs. Pare.

### 3.5. Cardiac METTL3 regulates the Akt/mTOR signaling pathway in irradiation

Thirty-six mouse heart samples were grouped by us according to high or low expression of METTL3, and 336 up-regulated differential genes were identified in the radiological dataset GSE218447. KEGG enrichment analysis was performed to select the top 10 pathways, and the results showed that more genes were significantly enriched in the PI3K-Akt pathway, which showed a strong correlation with METTL3 (Fig. S6A). Therefore, we hypothesized that the PI3K-Akt pathway plays an important role in the regulation of RIHD by METTL3.

Western blot analysis demonstrated a dose-dependent increase in the phosphorylation levels of mTOR and Akt in CFs (Fig. 5A). Subsequent experiments involving knockdown and overexpression of METTL3 in CFs revealed that the phosphorylation levels of mTOR and Akt decreased after METTL3 knockdown (Fig. S7A), whereas they increased after METTL3 overexpression (Fig. S7B). Additionally, the phosphorylation levels of mTOR and Akt decreased in irradiated CFs following transfection with shMETTL3 (Fig. 5B).

**Fig 5.**
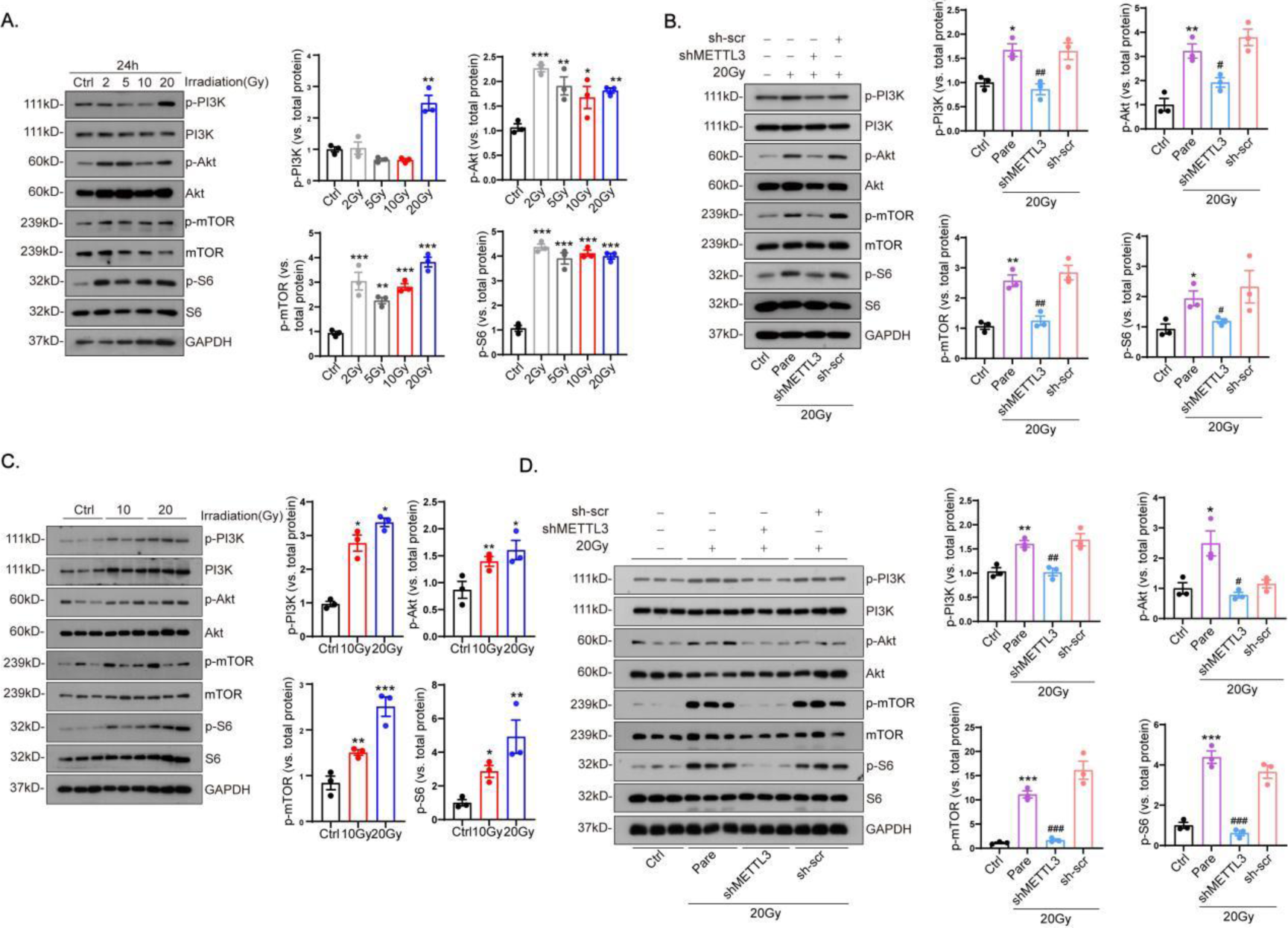
METTL3 initiates the Akt-mTOR cascades during irradiation Adult mice and cardiac fibroblasts were subjected to irradiation at doses of 0Gy (control), 2Gy, 5Gy, 20Gy (Pare), 20Gy with cardiac co-transfection of shMETTL3(shMETTL3) or sh-vector (sh-scr), with exposure for 4weeks and 24 hours, respectively. Phosphorylation of signaling molecules including PI3K, Akt, S6 and mTOR in heart tissues (A and B) and cardiac fibroblasts (C and D) were quantified using westering blotting. Data are shown as mean±SEM for groups of 3. The results were analyzed using one-way ANOVA with Tukey’s post hoc test and confirmed by three independent experiments. ***P<0.001, **P<0.01, *P<0.05 vs. Ctrl; ###P<0.001, ##P<0.01, #P<0.05 vs. Pare.

Mice were transfected with shMETTL3 or sh-scr viruses, and CFs were extracted from mouse hearts after irradiation at different doses. Immunofluorescence staining confirmed the presence of CFs, and subsequent western blot analysis confirmed enhanced phosphorylation levels of Akt and downstream signaling in cardiac CFs after irradiation (Fig. 5C). Conversely, in irradiated cardiac CFs transfected with shMETTL3 virus, the levels of Akt and downstream signaling phosphorylation were decreased compared to the former (Fig. 5D). These findings suggest that METTL3 knockdown, by inhibiting the Akt/mTOR signaling pathway and reducing myocardial fibrosis, significantly protects against cardiac injury resulting from post-irradiation fibrosis.

## 4. Discussion

This study demonstrates that X-ray radiation induces upregulation of METTL3, which is associated with reduced cardiac function and exacerbated cardiac fibrosis. Subsequent experiments revealed that METTL3 promotes cardiac fibroblast proliferation, migration, and fibrosis by activating the Akt/mTOR pathway. Additionally, our findings indicate that X-irradiation significantly elevates TGF-β1 levels, with METTL3 knockdown mitigating this effect and attenuating X-irradiation-induced fibrosis (Fig. 6).

**Fig 6.**
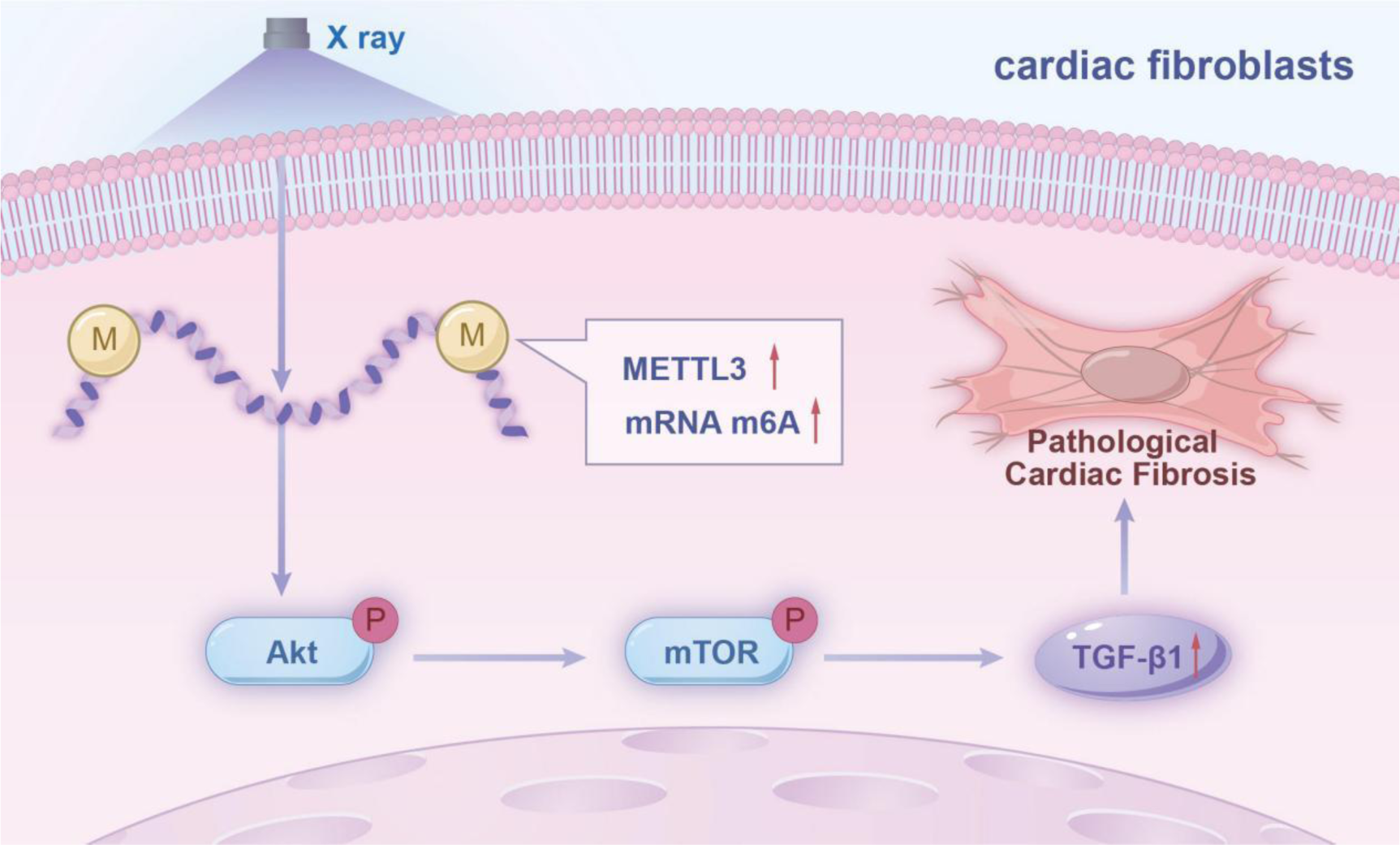
A model for the effect of m6A RNA methylation-regulated Akt/mTOR signaling pathway on radiation-treated cardiac fibroblasts.

Previous studies have shown that cardiac radiotherapy induces myocardial fibrosis at early stages, and enhances myocardial remodeling at later stages in mice^[14]^. In studies of RIHD, radiation doses reported in the literature include several Gy values^[16]^, but most are >10 Gy and up to 50 Gy^[17]^. Researchers found that increasing the radiation dose to 20 Gy led to cardiac fibrotic damage and structural damage^[14]^. Our current study confirms that 20 Gy radiation leads to cardiomyocyte hypertrophy and myocardial fibrosis in mice, in parallel with similar results in CFs. Myocardial fibrosis is a complex biological process involving many cellular activities and molecules. Cardiac fibroblasts, which make up 10–25% of the total number of cells in the myocardium have an essential role in maintaining structural and mechanical cardiac integrity^[18]^. CFs are responsible for maintaining the composition and organization of the extracellular matrix (ECM) in the cardiac wall, and they are central mediators of fibrosis. In response to injury, CFs are activated and acquire a pro-fibrotic phenotype commonly referred to as myofibroblasts, which is characterized by proliferative activity, excessive ECM production, and contractile function due to the expression of smooth muscle α-actin (α-SMA) ^[19]^.The activity of CFs is crucial in cardiac fibrosis because these cells are associated with tissue remodeling during fibrosis^[20]^. In addition, cardiac fibrosis is characterized by excessive proliferation of CFs. The transformation of fibroblasts into secretory, matrix-producing, and contractile cells (known as myofibroblasts) is a key cellular event in many fibrotic diseases. The transition from fibroblasts to myofibroblasts is a multifactorial process mediated by a number of factors^[21]^. Injured cardiomyocytes are also capable of producing fibrotic mediators in response to stress, thereby promoting fibroblast activation^[7]^. Our study corroborated these findings by demonstrating a dose-dependent upregulation of the myofibroblast marker α-SMA in CFs and mouse hearts post-irradiation This suggests enhanced fibroblast activation and increased extracellular matrix (ECM) proteins in irradiated, damaged hearts. These ECM proteins deposit in the interstitial matrix of the cardiac myocardium and induce to cardiac dysfunction.

Of the more than 100 chemically distinct coding and non-coding RNA modifications identified to date, N6-methyladenosine (m6A) is the most abundant and influential modification of mRNAs^[22]^. Modification of m6A in RNA transcript is a prevalent form of post-transcriptional modification of mRNAs, which has a significant impact on cardiac gene expression, cellular proliferation, and growth by regulating RNA stability and translation^[23]^. METTL3 was the first m6A methyltransferase to be identified. METTL3 is involved in numerous pathophysiological processes, including the cell cycle, apoptosis, and inflammation, as well as cellular behaviors including proliferation, migration, invasion, and differentiation^[24]^. Prabhu Mathiyalagan^[25]^ et al. demonstrated the critical role of the FTO-dependent cardiac m6A methylome in modulating heart failure and cardiac contraction. However, METTL3 has not been investigated in RIHD. Our study revealed that METTL3 expression was up-regulated in irradiated hearts, thereby intensifying cardiac fibrosis. Similar findings in irradiated CFs suggest a close association between METTL3 and RIHD. Our study also shows that knockdown of METTL3 led to recovery in cardiac function and reduction in fibrosis in both irradiated hearts and CFs, compared to irradiated controls, indicating that METTL3 negatively modulates RICF expression.

Akt is a serine/threonine protein kinase that serves as a central hub for signaling and regulates cellular functions, including survival, growth, and metabolism^[26]^. Studies have shown that Akt family members are expressed in the myocardium ^[27]^. Zhen-Guo Ma et al. discovered that cardioprotective effects of pipeline were negated by the overexpression of constitutively active Akt. Inhibition of AKT significantly attenuated pressure-overload-induced cardiac fibrosis^[28]^.Liu et al. discovered that CYP19A1-catalyzed estrogen biosynthesis promotes vascular abnormality through the upregulation of TGF-β via GPR30-AKT signaling^[29]^. Previous studies have shown that mTOR is a direct substrate for AKT kinase. mTOR is also a serine/threonine kinase, consisting of two complexes, mTORC1 and mTORC2^[30,31]^. Finckenberg et al.^[32]^ found that when the mTOR pathway was blocked with everolimus in double-transgenic rats, pro-fibrotic factors TGF-β1, Col I, and Col III expression levels were significantly down-regulated. Mitra et al.^[33]^ found that OSI-027, a dual mTOR inhibitor, inhibited TGF-β-induced expression of α-SMA, Col I, and Col III. AKT/mTOR may be involved in the regulation of cardiac fibrosis. Consistent with the above studies, we found that the fibrosis level of CFs increased after irradiation, and the phosphorylation levels of Akt, mTOR, and downstream S6 in CFs and cardiac tissues were up-regulated according the irradiation dose, while knockdown of METTL3 both in vivo and in vitro the levels of phosphorylation of mTOR and downstream S6 went down. This suggests that the upregulation of METTL3 after irradiation exacerbates cardiac fibrosis, potentially through the AKT-mTOR pathway.

TGF-β is a multifunctional cytokine that is released from fibroblasts at the site of myocardial tissue injury ^[34,35,36]^. TGF-β1, secreted by fibroblasts in the heart, has autocrine and paracrine functions, and continued self-induction of TGF-1 during repetitive injuries ^[37]^. TGF-β1-overexpressing mice exhibit significant cardiac hypertrophy with interstitial fibrosis^[38]^.Elevated levels of TGF-β1 in the blood and lungs have been documented during the early stages of radiotherapy^[39]^. Exposing human umbilical vein endothelial cells to radiation resulted in the development of a cell line model for sustained TGF-β1 release^[40]^. Elevated TGF-β levels prompt collagen synthesis, ultimately resulting in fibrosis^[41]^. The findings indicate that increasing of TGF-β1 in CFs and mouse hearts after irradiation is involved in the pathological process of myocardial fibrosis.

Our study has several limitations. First, it should be noted that the expression of METTL3 is subject to dynamic regulation in response to varying levels of irradiation. However, the specific mechanisms that govern this regulation are still not well understood. Second, the induction of METTL3 in disease models has been achieved using lentiviral vectors. However, the use of these vectors in clinical settings is limited because of the possibility for unintended effects on non-targeted areas, constraints in delivery methods, and concerns over safety. Third, we have discovered specific molecular targets that are influenced by METTL3 and play a role in regulating the phosphorylation of PI3K-Akt. However, it is important to thoroughly investigate additional molecules downstream that may possibly be involved in this progress.

In conclusion, Elevated expression of METTL3 is involved fibrotic remodeling of the heart after radiation, and METTL3 interacts with the Akt pathway to modulate the fibrotic response to pro-fibrotic stimuli. Based on the results of this study and other studies that have been talked about above, selectively targeting METTL3 silencing may be a new and effective way to stop heart failure and cardiac fibrosis caused by radiotherapy.

**Table 1.**
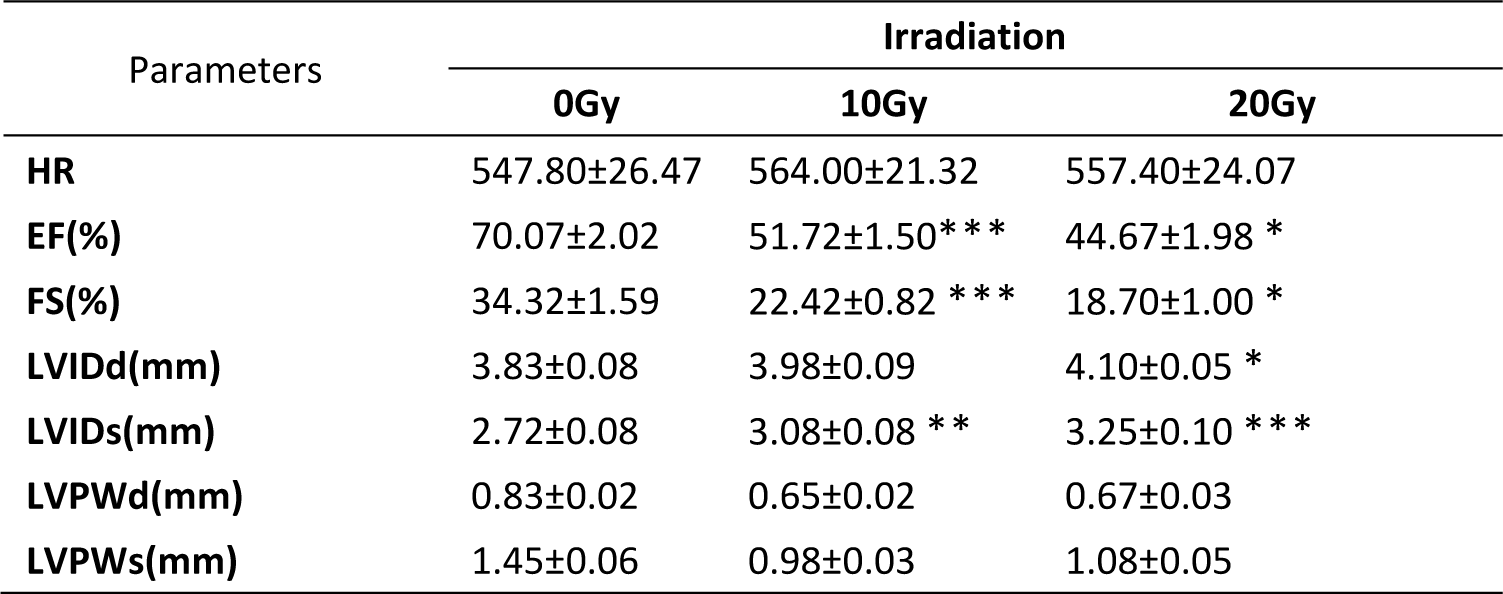
Radiation is associated with impaired heart function adult mice were subjected to irradiation for 4weeks. The heart function parameters were likewise assessed using echocardiography. (Heart function parameters including heart rate (HR), ejection fraction (EF); fraction of shortening (FS), left ventricular end-diastolic internal diameter (LVIDd), left ventricular end-systolic internal diameter (LVIDs), left ventricular end-systolic posterior wall thickness (LVPWs) and left ventricular end-systolic posterior wall thickness (LVPWs), were assessed using echocardiography. Data are expressed as mean ± SEM for groups of 6. All results were analyzed using one-way ANOVA with Tukey’s post hoc test and confirmed by three independent experiments. ***P<0.001,**P<0.01,*P<0.05 vs. Ctrl.

**Table 2.** Heart function of irradiation adult mice with cardiac co-transfection of shMETTL3(shMETTL3) or sh-vector (sh-scr) for 4 weeks. METTL3 is key to impaired heart function during irradiation adult mice were subjected to irradiation at doses of 0Gy (control), 20Gy (Pare), 20Gy with cardiac co-transfection of shMETTL3(shMETTL3) or sh-vector (sh-scr) for 4 weeks. Data are expressed as mean ± standard deviation for groups of 6. All results were analyzed using one-way ANOVA with Tukey’s post hoc test and confirmed by three independent experiments. ***P<0.001,**P<0.01,*P<0.05 vs. Ctrl.

| Parameters | Treatment |  |  |  |
| --- | --- | --- | --- | --- |
|  | Ctrl | Pare | shMETTL3 | sh-scr |
| **HR** | 542.00±26.81 | 582.00±24.99 | 567.00±21.46 | 565.6±20.97 |
| **EF(%)** | 68.67±1.75 | 37.97±1.14 *** | 59.20±3.37 *** | 36.77±1.64 |
| **FS(%)** | 33.32±1.29 | 15.40±0.55 *** | 27.05±2.09 *** | 14.92±0.76 |
| **LVIDd(mm)** | 4.13±0.23 | 4.28±0.28 | 4.15±0.11 | 4.90±0.11 |
| **LVIDs(mm)** | 3.70±0.34 | 3.65±0.23 * | 3.07±0.05 * | 4.17±0.13 |
| **LVPWd(mm)** | 0.75±0.02 | 0.70±0.04 | 0.67±0.03 | 0.68±0.05 |
| **LVPWs(mm)** | 1.18±0.05 | 0.97±0.08 | 1.00±0.07 | 0.97±0.10 |

## Non-standard Abbreviations and Acronyms

RIHD: radiation-induced heart disease
CFs: cardiac fibroblasts
ECM: extracellular matrix
m6A: N6-methyladenosine modification
METTL3: methyltransferase like 3
RICF: radiation-induced myocardial fibrosis
TGF-β1: transforming growth factor β1
IR: ionizing radiation
SR: Sirius Red
WGA: wheat germ agglutinin
α-SMA: smooth muscle α-actin

## Acknowledgments

This study was funded by the Department of Cardiology of the Second Affiliated Hospital of Soochow University. We thank all the staff for their help with data collection.

## Author Contributions

XSG and ZJ conceived the study and designed the study protocol. QSS and ZJJ constructed animal model and collected sample. TC and ZJ performed the echocardiographic analyses. ZJJ and XL conducted the literature review and statistical analysis. QSS and XL drafted the manuscript. ZJJ and XSG reviewed the manuscript for intellectual content, made revisions as needed. All authors contributed to editorial changes in the manuscript and read and approved the final manuscript.

## Perspectives

**Competency in Medical Knowledge**: METTL3 is a new target for treating post-radiotherapy myocardial fibrosis, regulating cardiac fibroblast activation via activation of the Akt pathway in IR mouse models.

**Translational Outlook:** This study provides additional evidence for METTL3 promoting RICF. Further clinical studies are needed to assess the potential adverse effects of radiation-induced METTL3 upregulation on cardiovascular diseases.

